# Effect of Aqueous Garlic Extract on Hyperlipidaemia in Rats: A Randomised Case-Control Study

**DOI:** 10.1101/2023.08.10.552555

**Authors:** Gamel Sankarl, Enoch Akyeampong, Micheal Sarhene, Rexford Kporvie Anai

**Author notes:** **Corresponding Author** Gamel Sankarl Department of Molecular Medicine School of Medical Sciences Kwame Nkrumah University of Science and Technology.

## Abstract

This study aimed to determine the effect of garlic extract on hyperlipidemia in rats. This was a randomised case-control study in 42 rats comprising seven groups with six animals per group. Group I consisted of rats fed with 100% Normal rat chow serving as control, Group II: Butter diet, Group III: Butter diet + Atorvastatin, Group IV: Butter diet + Garlic extract, Group V: rats injected with dexamethasone, Group VI: Dexamethasone + Atorvastatin, Group VII: Dexamethasone + Garlic extract. Blood was collected from all the animals and the separated serum was subjected to the estimations of lipoproteins levels and coronary risk. There was a significant increase in the concentration of TC in the butter control group (BCG) compared to the concentration of the LDL-C (p=0.028) of the normal control group (NCG). The concentration of TG was significantly higher in DCG when compared with TG concentration in NCG (p<0.0001) and BCG (p=0.0002). Total Cholesterol level was significantly higher in BCG when compared to BGG (p<0.0001), BAG (p=0.0001) and NCG (p=0.023) which typifies hypercholesterolemia. Hyperlipidemia was induced by the long-term high-fat diet but lipid levels were reduced after the administration of aqueous garlic extract and atorvastatin.

## INTRODUCTION

Garlic (*Allium sativum*) has become the toast of modern medical research because it showed significant hypolipidaemic effects in clinical and experimental research studies [1]. Garlic has reported that many people have been reluctant to use a lot of human health benefits due to its invaluable biological effects and these include being; an antimicrobial agent, an antioxidant, an antidiabetic agent, a risk factor reducing agent for cardiovascular diseases, an immune system stimulant, a fibrinolytic agent, an agent for hepatoprotection and have enhanced abilities in the detoxification of foreign compounds [2].

One of the prominent sulphur-containing compounds of garlic is ‘allin’ which is always inactive until the garlic bulb is crushed. When allin is crushed, it becomes activated under the catalytic action of the enzyme allinase and converts into ‘allicin’; an active ingredient involved in the inhibition of HMG-CoA reductase and other principal enzymes responsible in the biosynthesis of cholesterol [3]. Some researchers have reported that many people have been reluctant in the usage of garlic because they do not enjoy its peculiar smell [4-6].

According to Steiner and Li [7], garlic have very effective preventive properties thatcause the prevention of a number of heart related diseases especially dyslipidaemias and hypertension. Garlic is also known to exert cholesterol and triglyceride lowering effects [8, 9]. It is antithrombotic [10], antimicrobial [11], antihypertensive [12, 13] and has great anti-hyperlipidemic effects [14, 15]. Additionally, garlic possesses some potent cardio-protective properties that enables it to effectively increase good cholesterol (high density lipoprotein-HDL) whilst concurrently decreasing the concentrations of bad cholesterol (low-density lipoprotein-LDL) and total cholesterol (TC) [2, 4].

Hyperlipidaemia has been implicated as one of the primary precursors for the development of cardiovascular diseases (CVDs) and CVDs have been reported to be responsible for an estimated 17.5 million deaths [16]. CVDs have also been reported by Livesay [17], to have caused 9.2% of deaths in Africa. Sadly, CVDs constitute one of top two leading causes of deaths in Ghana [16]. A research study by Bosu [18] also indicated that a higher percentage (14.5%) of Ghanaians die as a result of CVDs than from malaria which is 13.4%. Sanuade, Anarf and Aikins [19] reported from their 5-year (2006-2010) review of autopsies carried out at the Pathology Department of the Korle-Bu Teaching Hospital that 22.5% of deaths in Ghana result from CVDs.

These research reports make it very imperative to correct hyperlipidaemia since it has become a silent killer and for that matter, a condition of major medical concern needing immediate attention. In an attempt to treat hyperlipidaemia and other cardiovascular related diseases, physicians and clinicians in general have resorted to the use of statins. Statins however could have severe side effects [20-22] and hence have led to patient’s non-compliance. Besides, statins are expensive chemical therapeutic products that are not easily accessible to poor patients. Given this background, if garlic (a natural herbal product) which is relatively affordable and accessible is proven to be an effective agent against hyperlipidaemia then it will be a preferred alternative treatment option for hyperlipidaemia. It is in the light of this development that this study, aimed at evaluating the effect of aqueous garlic extract on hyperlipidaemia, has become necessary. The findings of this research study will help establish the efficacy of garlic extract against hyperlipidaemia. It will help confirm whether garlic should or should not be used as an alternative treatment option for hyperlipidaemia.

## MATERIALS AND METHODS

### Study design

The study was a randomized case-control which used animal models as study subjects at the animal house of the Department of Pharmacology, Kwame Nkrumah University of Science and Technology.

All methods of the study were carried out in accordance with Declaration of Hesinkin and Data Protection Regulation of Ghana, thus, Data Protection Act 2012 (Act 843). Written consent form was applicable in this study.

### Selection of experimental rats

Rats were selected using the non-probability convenience sampling technique to group 42 experimental albino rats into seven groups of six rats per group.

### Inclusion Criteria

Rats with body weight between 200-250 grams were selected for the study. Only healthy rats with normal lipid profile indices were selected for the study.

### Exclusion Criteria

Rats with abnormal lipid profile indices were exempted from the study. Rats with body weights less than 200-250 grams were not selected.

### Sample size of experimental rats

Forty-two (42) male albino rats were obtained from Korle-Bu Animal House for the research. These rats were eight weeks old and weighed between 200-250 grams as at the start of the study.

### Preparation of Animals for Study

The rats were housed in steel cages and maintained under conditions of 12 hours light and 12hours dark cycle at an optimum temperature. These animals were divided into seven groups with six animals per group. The grouping of the animals was as follows:

Group I: Control (100% Normal rat chow) for 35 days.

Group II: Butter diet (70% Normal diet + 30% butter) for 35 days.

Group III: Butter diet + Atorvastatin (10 mg/kg body weight) for 35 days.

Group IV: Butter diet + Garlic extract (200mg/kg body weight) for 35 days.

Group V: Dexamethasone (100% Normal diet + 10mg/kg body weight) for 5days.

Group VI: Dexamethasone + Atorvastatin (10mg/kg body weight) for 5 days.

Group VII: Dexamethasone + Garlic extract (400mg/kg body weight) for 5 days.

As indicated in the various groups, the animals received different treatments. They were broadly divided into three groups comprising of;

1. The Normal Control Group [I (NCG)]
2. The Diet or Butter Groups [II (BCG), III (BAG) and IV (BGG)]
3. The Drug or Dexamethasone Groups [V (DCG), VI (DAG) and VII (DGG)]

With the exception of the rats in the Normal Control group (Group I) which received normal rat chow and tap water for 35 days, all the other rats received diets or drugs that induced hyperlipidaemia in them.

### Inducing hyperlipidaemia in experimental rats

To induce hyperlipidaemia in the rats for the study, high fat-diet and drug were given to the rats in the remaining groups. The butter groups (BCG, BAG and BGG) received butter diet. This high fat-diet was prepared using a saturated fat butter brand called Cook Brand and normal rat chow. 30 grams of the butter was added to 70 grams of the normal rat chow and stirred to form a uniformly mixed paste (soft cake). This was then served to the rats on a daily basis. A fresh feed was prepared for the rats every day for 35 days.

Rats of Group II served as the Butter Control Group (BCG) and received only the butter diet and tap water for 35 days. Groups III (BAG) and IV (BGG) also received the butter diet for the first 35 days and respectively received atorvastatin and garlic extracts for another 35 days after the induction. Another way hyperlipidaemia was induced in the rats for this study was through the use of the drug dexamethasone. The dexamethasone was bought from a licensed Pharmaceutical shop called Western Pharmacy. The right dosage calculations were made before administering the dexamethasone on the rats. The drug administration was done using a syringe. The administration was done through subcutaneous injection for 5 days.

Rats of Group V served as the Dexamethasone Control Group (DCG) and received only the normal rats chow and tap water for 5 days. They also received a subcutaneous administration of 10mg/kg body weight of the dexamethasone. Rats of Groups VI (DAG) and VII (DGG) also received 10mg/kg body weight subcutaneous administration of dexamethasone plus normal rat chow and tap water. However, they also respectively received Atorvastatin and Garlic Extract concurrently. With the exception of the garlic which was done using an oral gavage, the dexamethasone and atorvastatin administrations were done through subcutaneous injections for 5 days.

### Treating induced-hyperlipidaemia in rats

After the hyperlipidaemia induction was successfully done, it was now expedient to study how aqueous garlic extract affected hyperlipidaemia. To achieve this task, atorvastatin; a statin drug was used as a positive control drug whilst the aqueous garlic extract was used as a negative control drug. These two were prepared and administered to the rats at different time intervals depending on the group.

### Administering Atorvastatin to Hyperlipidaemic Rats

The Atorvastatin tablets were bought from a local pharmaceutical store. The tablets were crushed using a pestle and mortar into powder. The powdered Atorvastatin was then weighed with a sensitive weighing balance (Sartorius-TE 214S). The weighted amount of 2mg was then dissolved in 10ml of water to form a stock solution. This stock solution of Atorvastatin was then administered to the rats at 10mg/kg body weight. This was given to them orally through the use of an intra-gastric tube.

The Atorvastatin was administered to the rats of Groups (III and VI). Thus, each rat in the butter diet group; Group III (BAG) received 10mg/kg body weight of the Atorvastatin for 35 days whilst each rat in the dexamethasone drug group; Group VI (DAG) also received 10mg/kg body weight of Atorvastatin orally for 5 days.

### Acute Toxicity Studies

The experimental rats were randomly selected and subjected to an acute toxicity study before a required extract dosage was administered. Prior to the extract administration, the rats were fasted overnight. They only received water during the fast period. The intra-gastric administration of the extract was done starting at a dose of 10mg/kg body weight for 14 days. If for a particular dose, mortality was observed in two rats, the exact dosewas repeated in two other rats separately to be sure of its toxicity. In the event that mortality was not observed, the method of administration was repeated with each new administration having a higher dose range such as from 25, 50, 100, 200, 400 and 1000mg/kg body weight [23].

### The Administration of Garlic Extract to Hyperlipidaemic Rats

The garlic bulbs were bought from a local market within the Kumasi metropolis. The garlic bulbs were peeled with a knife. And the aqueous garlic extract was prepared using a modified method of Martha et al. [24]. Thus, 50 grams of the peeled garlic was weighed using a weighing scale. The weighted garlic was homogenized by blending it in 100ml of cold distilled water. The homogenized mixture was then filtered trice through a layered folded gauze. The mixture was then centrifuged using the Mikro 220R [Hettich Zentrifugen] centrifuge at 5000rpm for 10 minutes. The clear supernatant was then collected into a sterile bottle and used for the administration. Each bottle of the aqueous garlic extract was refrigerated at an optimal temperature for a week. A fresh aqueous garlic extract was prepared every other week for administration. The aqueous garlic extract was administered to rats of Groups (IV and VII). Thus, the butter diet, group IV (BGG) and the dexamethasone drug, group VII (DGG).

For the rats of Group IV (BGG), the aqueous garlic extract was administered for 35 days. This was done after the initial 35days of the hyperlipidaemia induction using butter diet. The aqueous garlic extract was administered using an intra-gastric tube. Each rat received 200mg/kg body weight of the aqueous garlic extract daily for the entire 35days. Each rat of the dexamethasone group; Group VII (DGG) were given 400mg/kg body weight of the aqueous garlic extract using an intra-gastric tube as well. However, this was done concurrently with the administration of the dexamethasone for 5 days.

### Measuring the body weights of experimental rats

At the initial stage of the study, the body weights of the rats were taken using a digital weighing scale. Their weights were then taken in the subsequent days or weeks all through the study period. The body weights of the rats of Group I (the Normal Control Group) was taken on a weekly basis for 35 days. Thus, on Day(s): 0, 7, 14, 21, 28 and 35. It was done over a five-week period. During the hyperlipidaemia induction, the body weights of the rats of Group II (Butter Control Group) were taken over a five-week period. Thus, on days 0, 7, 14, 21, 28 and 35. During the hyperlipidaemia reduction (atorvastatin and garlic extract administration), the rats of Group III (Butter + Atorvastatin Group) and Group IV (Butter + Garlic Group) were taken on a weekly basis for five weeks. Thus, the body weights were measured on days 0, 7, 14, 21, 28 and 35. The average body weights of the rats in Group II (Butter Control Group) were then recorded to determine the effect of the butter diet on the weight of the rats during the hyperlipidaemia induction period.

Also, the average body weights of the rats of Groups III (BAG) and IV (BGG) were recorded during the hyperlipidaemia induction to determine the effect of butter diet on the rats. After which the average body weights were monitored and recorded during the hyperlipidaemia reduction to respectively determine the effects of atorvastatin and aqueous garlic extract on the body weights of the rats of Groups III (BAG) and IV (BGG). The body weights of the rats of Group V (Dexamethasone Control Group), Group VI (Dexamethasone + Atorvastatin) and Group VII (Dexamethasone + Garlic) were taken on a daily basis for 5 days. This was done on days 0, 1, 2, 3, 4 and 5. And it was done simultaneously as the hyperlipidaemia induction and the hyperlipidaemia reduction was concurrent.

The average bodyweights of the rats in Groups V (DCG), VI (DAG) and VII (DGG) were recorded to respectively determine the effects of dexamethasone, dexamethasone + atorvastatin and dexamethasone + garlic aqueous on the body weights of the experimental rats.

### Biochemical Analysis of Collected Blood Samples

After both the processes of hyperlipidaemia induction and the administration of atorvastatin and garlic extract, the rats were sacrificed and the blood samples were collected for analysis. Prior to taking the blood samples for lipid profile analysis, the rats were fasted for 12hours. Blood samples of Group I (NCG) rats were taken after 35 days of receiving normal rat chow and normal tap water. The rats were sacrificed through decapitation and 2ml of the blood sample was collected into gel separator tubes. The blood samples were then processed and well-labelled before the biochemical analysis.

Blood samples of rats of Groups II, III and IV (the Butter Groups) were taken at different time intervals using the decapitation method. The blood samples of Group II (Butter Control Group) was taken after the 35 days of hyperlipidaemia induction using butter diet. The blood samples of Group III (Butter +Atorvastatin) and Group IV (Butter + Garlic Extract) were taken after the hyperlipidaemic rats were respectively administered with atorvastatin and garlic extract. Thus on the 70^th^ day. After the first 35days of the hyperlipidaemia induction, the rats in both groups (Group III and Group IV) respectively received Atorvastatin and Garlic Extract for the next 35 days. The rats were sacrificed through decapitation and the blood samples collected into gel separator tubes. The blood samples of rats of Group V (Dexamethasone Control Group) were taken 5days after hyperlipidaemia induction. Blood samples of rats of Group VI (Dexamethasone + Atorvastatin) and Group VII (Dexamethasone + Garlic Extract) were also taken 5days after atorvastatin and garlic extract administration.

All blood samples were collected in the morning (after an overnight fast) between the hours of 8 and 9am. The blood samples were then processed, labelled and were made ready for the biochemical analysis. The biochemical analysis; lipid profile analysis was done using the Automated Junior Chemistry Analyser. Prior to analysing in the Chemistry Analyser, the blood samples in the tubes were spun in a centrifuge at 3000 x g for 10 minutes to separate the plasma from the sera. The lipid sample was then stored in a refrigerator at a temperature of -80 °C until the analysis. The results from the analysis provided the concentrations of the lipid profile parameters; total cholesterol (TC), triglycerides (TG), high density lipoproteins (HDL) and very low density lipoproteins (VLDL) in their right indices. These indices helped in elucidating the anti-hyperlipidaemic effect of aqueous garlic extract.

### Biochemical Variables

The biochemical variables considered for this study are the lipid profile parameters namely; total cholesterol, high-density lipoprotein, low-density lipoprotein and triglycerides. To measure these lipid profile parameters, aliquots were dispensed into cryotubes after thawing. The tubes with the sera were then placed at pre-programmed positions in the autoanalyser and the analysis was done in batches. All the reagents used in the analysis were made by the same manufacturer. LDL-cholesterol was calculated according to Friedewald equation [25].

LDL-cholesterol = (Total cholesterol) – (HDL-cholesterol) – (Triglycerides/2.17).

### Total Cholesterol (TC)

The biochemical analytical procedure for total cholesterol was first recorded and described by Trinder, (1969). The procedure postulates that, cholesterol undergoes a catalytic reaction when exposed to cholesterol esterase. This causes its breakdown resulting in the production of particulate cholesterol and fatty acids. There is a hydrolysis of its esters which is facilitated by a fungal lipase leading to the release of hydrogen peroxide which in turn produces the oxidative coupling of phenol with 4 – aminophenazone (4 AP). The catalytic reaction process is made possible by peroxidase and a red quinoneimine is formed. The red quinoneimine (dye) has a maximum absorption rate of 500-505nm. The intensity of the red colour produced is directly proportional to the total cholesterol read.

### High-Density Lipoprotein Cholesterol (HDL-C)

The method is based on a modified polyvinyll sulfonic acid (PVS) and polyethylene-glycol methyl ether (PEGME) coupled classic precipitation method with the improvements in using optimized quantities of PVS/PEGME and selected detergents [26]. LDL, VLDL and chylomicrons react with PVS and PEGME and the reaction results in inaccessibility of LDL, VLDL and chylomicrons by cholesterol oxidase and cholesterol esterase. The enzymes selectively react with HDL to produce hydrogen peroxide through a Trinder reaction in the presence of N, N-Bis(4-sulfobutyl)-3-methylaniline (TODB). The intensity of the red colour produced is directly proportional to the HDL-cholesterol in the sample when read at 560 nm.

### Triglycerides (TG)

The method for the analysis is a modification of that of Trinder, (1969). Triglycerides in the sample are hydrolyzed by lipase to glycerol and fatty acids. The glycerol is then phosphorylated by adenosine-5-triphosphate to glycerol-3-phosphate and adenosine-5-diphosphate in a reaction catalyzed by glycerol kinase (GK). Glycerol-3-phosphate is then converted to dihydroxyacetone phosphate (DAP) and hydrogen peroxide by glycerophosphate oxidase. The hydrogen peroxide then reacts with 4-aminoantipyrine (4-AAP) and 3-5 dichloro-2-hydroxybenzene (3, 5-DHBS) in a reaction catalyzed by peroxidase (POD) to yield a red coloured quinoneimine dye in the sample. The intensity of the color produced is directly proportional to the concentration of Triglycerides in the sample.

### Statistical Analysis

The data obtained from the research study was analysed using SPSS version 21 statistical software. The data was subjected to analysis of variance. Results obtained from the analysis are expressed as mean ± standard error of mean (SEM). To assess the statistical significance, One-Way Analysis of Variance was used. This was followed by the analysis of mean values using the Dunnett’s comparison test which helped in determining whether or not there were any significant differences existing between the groups. A P-Value of less than 0.05, thus, (P<0.05) was considered statistically significant.

## RESULTS

### Effect of Butter and Dexamethasone on the Lipid Profile of Rats

The administration of 30% butter enriched diet and 10mg/kg body weight of dexamethasone to experimental rats showed changes in the serum lipid and lipoprotein levels of these rats. The butter caused significant increase in the concentration level of TC in the Butter Control Group (BCG) compared to the concentration levels of the LDL-C (3.08±0.18 vs. 2.33±0.22, p=0.028) of the Normal Control Group (NCG). (Table 1).

**Table 1.** Effect of Butter and Dexamethasone on the Lipid Profile of Rats.

| Variables | NCG (A) | BCG (B) | DCG (C) | p-value | Significant pairs |
| --- | --- | --- | --- | --- | --- |
| **TC (mmol/L)** | 2.33±0.22 | 3.08±0.18 | 2.65±0.13 | **0.035** | A vs. B (0.028) |
| **HDL (mmol/L)** | 0.95±0.06 | 1.00±0.08 | 0.88±0.13 | 0.634 | 0.634 |
| **LDL (mmol/L)** | 1.23±0.11 | 1.52±0.15 | 0.83±0.18 | **0.022** | A vs. C (**0.017**) |
| **TG (mmol/L)** | 0.35±0.07 | 0.53±0.04 | 0.63±0.33 | **<0.0001** | A vs. C (**<0.0001**), B vs. C (**0.0002**) |
| **CR** | 2.40±0.12 | 3.10±0.28 | 3.38±0.50 | 0.145 |  |

The values are expressed as mean ± SEM, (n=6), both groups (BC and DC) were compared with the Normal Control Group, TC: Total cholesterol, TG: Triglyceride, LDL: Low-density lipoprotein, HDL: High-density lipoproteins, SEM: Standard error of mean, NCG: Normal Control Group, BCG: Butter Control Group, DCG: Dexamethasone Control Group.

Also, there were changes in the concentrations of the lipid profiles of the experimental rats in the dexamethasone group (ie. Control, atorvastatin and garlic groups). (Table 1). Statistically, there were no significant (p=0.145) differences between the calculated coronary risk ratios of the NCG (2.40±0.12), BCG (3.1±0.28) and the DCG (3.38±0.50). These values clearly depict that the BCG and the DCG had developed hyperlipidaemia and therefore have higher risk of developing coronary heart conditions (CHC) compared tothe NCG. (Table 1)

### Effect of Garlic Extract on High-Density Lipoprotein Cholesterol (HDL-C)

The concentration of HDL-C in the Butter Garlic Group (BGG) (0.98±0.06mmol/L) ofthe butter-induced hyperlipidaemia group showed an insignificant increase in concentration upon the administration of aqueous garlic extract when compared to the Normal Control Group (NCG) (0.95±0.06mmol/L). (Table 2). The LDL-C level of BGG was significantly (p=0.002) lower compared to that of the BCG. Total Cholesterol (TC) level was significantly higher in BCG when compared to BGG (3.08±1.16 vs. 1.75±0.76mmol/L, p<0.0001), BAG (3.08±1.16 vs.1.8±0.16mmol/L, p=0.0001) and NCG (3.08±1.16 vs.2.33±0.22mmol/L, p=0.023) which typifies hypercholesterolaemia. (Table 2). The values also indicate that aqueous garlic extract have greater reducing-effect on TC than the positive control drug, Atorvastatin. Triglycerides (TG) values of BGG (p=0.0008) and BAG (p=0.001) were significantly higher than the value of NCG (0.35±0.72mmol/L). These values however indicated that aqueous garlic extract and atorvastatin exerted hypolipidaemic or anti-hyperlipidaemic effects when compared to the BCG and the NCG. The coronary risk ratio of the butter-induced hyperlipidaemic group indicated a statistically significant increase in value of the butter control group (BCG) when compared to the normal control (NCG) value of (3.10±0.28 vs. 2.40±0.30, p=0.014). (Table 2).

**Table 2.** Effect of Garlic Extract on Butter-Induced Hyperlipidaemia.

| Variables | NCG (A) | BCG (B) | BAG (C) | BGG (D) | p-value | Significant pairs |
| --- | --- | --- | --- | --- | --- | --- |
| **TC (mmol/L)** | 2.33±0.22 | 3.08±0.18 | 1.80±0.38 | 1.75±0.16 | **<0.0001** | A vs. B (**0.023**), B vs. C (**0.0001**), B vs. D (**<0.0001**) |
| **HDL (mmol/L)** | 0.95±0.06 | 1.00±0.08 | 1.02±0.14 | 0.98±0.10 | 0.637 |  |
| **LDL (mmol/L)** | 1.23±0.11 | 1.52±0.15 | 0.85±0.09 | 0.83±0.08 | **0.0005** | B vs. C (**0.013**), B vs. D (**0.002**) |
| **TG (mmol/L)** | 0.35±0.07 | 0.53±0.04 | 0.68±0.05 | 0.67±0.03 | **0.0005** | A vs. C(**0.0008**), A vs. D (**0.001**) |
| **CR** | 2.40±0.12 | 3.10±0.28 | 1.93±0.25 | 2.20±0.27 | **0.018** | A vs. B (**0.014**) |

The values are expressed as mean ± SEM, (n=6). Both groups (BC and DC) were compared with the Normal Control Group, TC: Total cholesterol, TG: Triglyceride, LDL: Low-density lipoprotein, HDL: High-density lipoproteins, SEM: Standard error of mean, NCG: Normal Control Group, BCG: Butter Control Group, DCG: Dexamethasone Control Group

### Effect of Garlic Extract on High-Density Lipoprotein Cholesterol (HDL-C)

In the dexamethasone-induced hyperlipidaemia group, HDL-C concentration decreased in the DGG with a concentration of (0.90±0.06mmol/L) as compared to the NCG concentration of HDL-C (0.95±0.06mmol/L). (Table 3). Comparing its concentration level to that of the NCG and Dexamethasone Atorvastatin Group, it is a good indication that garlic extract raised the HDL-C which is very crucial in the treatment of hyperlipidaemia. Triglyceride levels of DGG in the dexamethasone-induced hyperlipidaemia group, showed a significant decrease as compared to its levels in the DCG (0.63±0.14 vs. 2.25±0.38mmol/L, p=0.001) and DAG (0.63±0.14 vs. 1.53±0.07mmol/L, p=0.031). Except for the reduced TG level of the NCG, this development implicates garlic extract as an effective agent against triglyceridaemia. (Table 4).

**Table 3.** Effect of Garlic Extract on Dexamethasone-Induced Hyperlipidaemia.

| Variables | NCG (A) | DCG (B) | DAG (C ) | DGG (D) | p-value | Significant pairs |
| --- | --- | --- | --- | --- | --- | --- |
| TC (mmol/L) | 2.33±0.22 | 2.65±0.13 | 2.13±0.21 | 1.53±0.20 | **0.005** | A vs. D (0.041), B vs. D (0.003) |
| HDL (mmol/L) | 0.95±0.06 | 0.88±0.13 | 0.72±0.05 | 0.90±0.06 | 0.111 |  |
| LDL (mmol/L) | 1.23±0.11 | 0.83±0.18 | 1.25±0.07 | 0.58±0.03 | **0.004** | A vs. D (0.003), C vs. D (0.04) |
| TG (mmol/L) | 0.35±0.07 | 0.63±0.33 | 1.53±0.07 | 0.33±0.18 | **<0.0001** | A vs. C (0.004), B vs. D (0.0001), A vs. B (p<0.0001) |
| CR | 2.40±0.12 | 3.38±0.50 | 2.97±0.10 | 2.21±0.20 | **0.033** | B vs. D (0.031), B vs. D (0.040) |

The values are expressed as mean±SEM, (n=6).TC: Total cholesterol, TG: Triglyceride, LDL: Low-density lipoprotein, HDL: High-density lipoproteins, SEM: Standard error of mean, DCG: Dexamethasone Control Group, DAG: Dexamethasone-Atorvastatin Group, DGG: Dexamethasone-Garlic Group. Both groups (DAG and DGG) were compared with the Dexamethasone Control Group and the Normal Control Group.

**Table 4:** Percentage (%) effect of mean difference of body weight of rats before and after administration of atorvastatin and garlic extract.

| Variables | Atorvastatin (Mean ± SEM) | Garlic Extract (Mean ± SEM) |
| --- | --- | --- |
| Before | 261.2±16.05 | 246.3±12.2 |
| After | 251.8±14.3 | 273.5±12.29 |
| p-value | 0.551 | **0.012** |
| Mean difference | -9.33 | 27.17 |
| % Mean Effect | 3.57% | 11.03% |

The values are expressed as mean ± SEM, SEM: Standard Error of Mean

The coronary risk ratio of the DGG was lower when compared to the Dexamethasone Control Group (3.38±0.50) and Dexamethasone Atorvastatin Group (2.97±0.10). (Table 3). The ratio implies that garlic extract has lipid-lowering protective properties and can therefore act as an anti-hyperlipidaemic agent against coronary heart related diseases.

The mean body weight of rats in the normal control group significantly decreased from week 1 to week 2 (207±16.75 vs. 193.0±14.78, p=0.043). Among the butter control group, mean body weight significantly (p<0.0001) increased from week 1 (185.0±17.42), week 2 (207.8±18.08), week 3 (237.2±17.30), week 4 (248.3±15.13) and week 5 (261.2±16.05). (Figure 1). The mean weight of the rats decreased from week 1 to week 2 after atorvastatin administration (261.2±16.05 vs. 248.2±14.25). No statistically significant differences were observed (p=0.456). (Figure 2). After the administration of garlic extract to the rats, no statistical significance was observed. However, the result showed a significant increase from week 4 to week 5 (257.2±13.39 vs. 273.5±12.29, p=0.048). (Figure 3)

**Figure 1:**
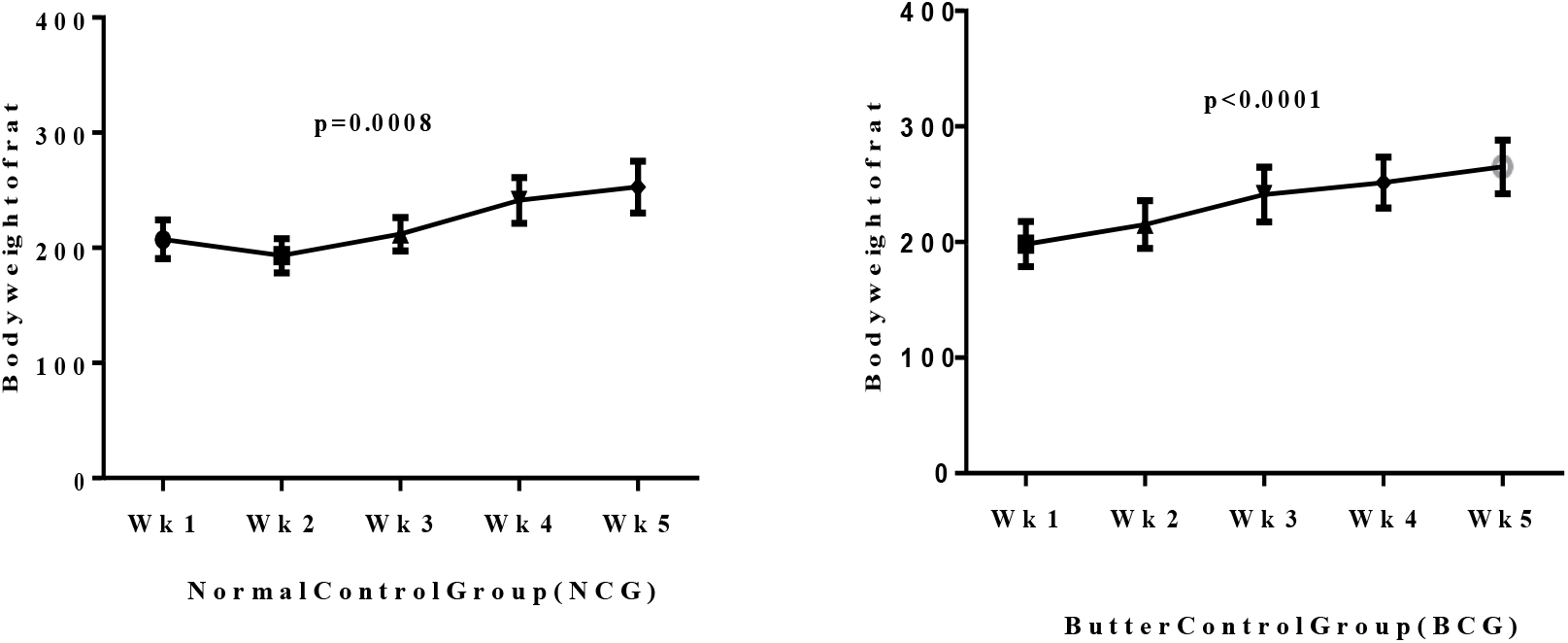
Body weight of normal control group and butter control group for 35days.

**Figure 2:**
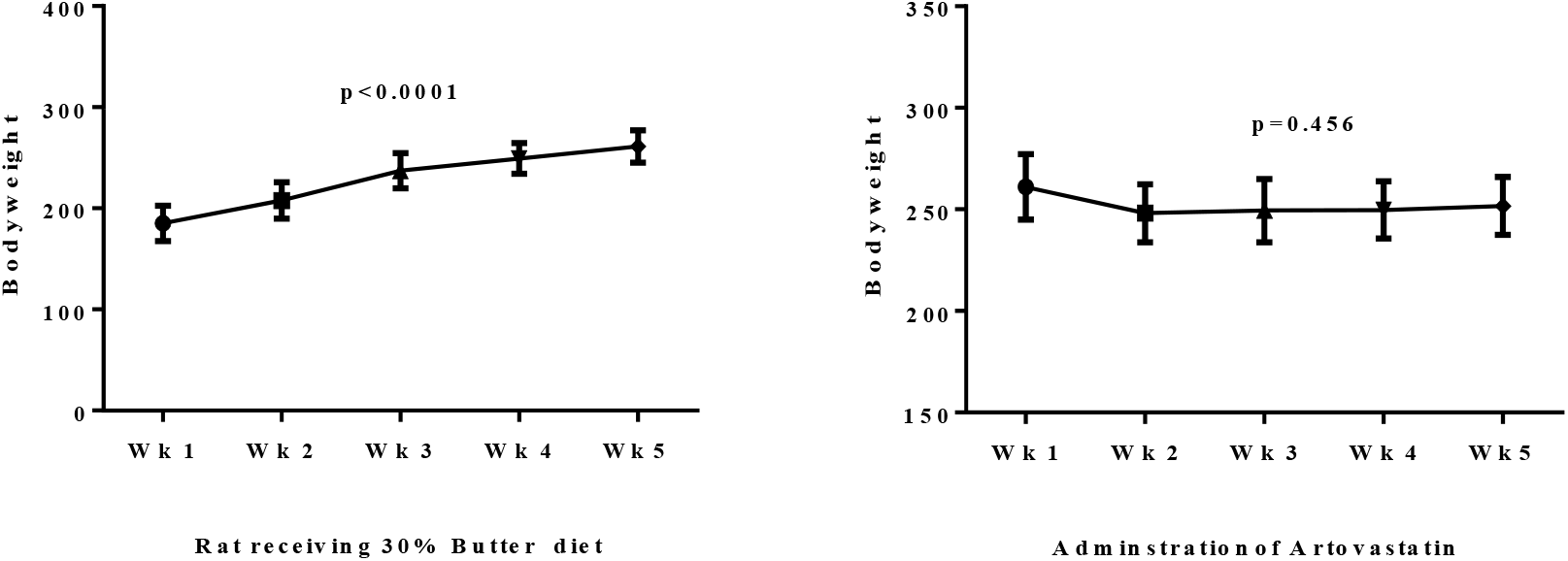
Body weight of rats after atorvastatin administration in 35 days.

**Figure 3:**
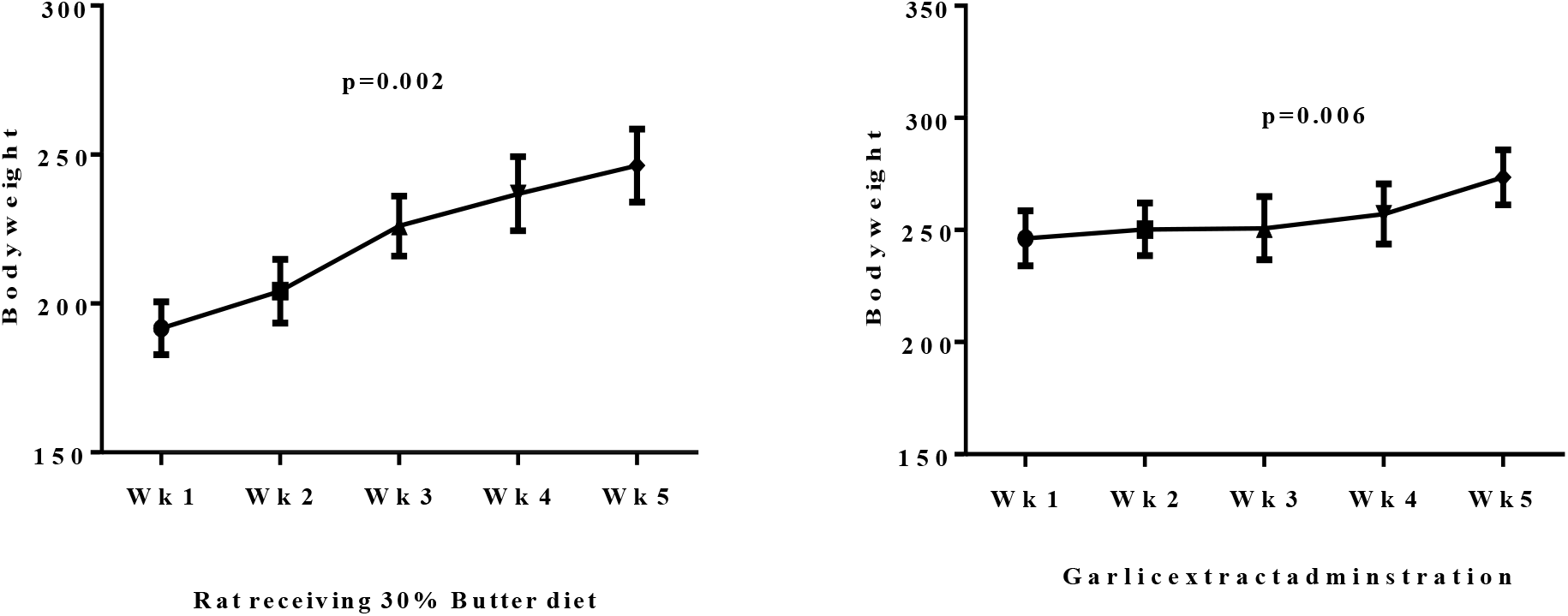
Body weight of rats after garlic extracts administration in 35 days.

Table 4 shows the percentage effect of mean difference of body weight of rats before and after administration of atorvastatin and garlic extract. The proportional effect of body weight of rats observed after atorvastatin administration was 11.03% while 3.57% was also observed after garlic administration. Also, percentage effect of mean difference (as shown in Table 5) after dexamethasone administration was 3.93%, while 5.46% was recorded after dexamethasone-garlic extract. As shown in Table 6, higher proportional effect of TC (41.6%), LDL-C (47.4%), coronary risk (37.7%), TG (28.3%) and HDL-C (2.0%) was observed after atorvastatin administration. Similar results were observed for LDL-C (44.7%), TC (43.2%), coronary risk (29.0%), TG (16.7%) and HDL-C (24.4%) respectively. Higher proportional effect of TG (32.0%), LDL-C (26.4%) TC (19.6%) HDL-C (19.3%) and CR (12.4%) was observed after dexamethasone-atorvastatin administration. (Table 7).

**Table 5:** Percentage effect of mean difference of body weight of rats before and after administration of dexamethasone, dexamethasone-atorvastatin, and dexamethasone-garlic extract.

| Variables | Dexamethasone (Mean ± SEM) | Dexamethasone-Atorvastatin (Mean ± SEM) | Dexamethasone-Garlic Extract (Mean ± SEM) |
| --- | --- | --- | --- |
| Before | 178.3±7.03 | 226.5±15.67 | 241.0±16.40 |
| After | 171.3±15.10 | 219.7±14.05 | 227.8±17.06 |
| p-value | 0.062 | 0.089 | **0.003** |
| **Mean difference** | -7.00 | -6.83 | -13.17 |
| **% Mean Effect** | 3.93% | 3.01% | 5.46% |

The values are expressed as mean ± SEM, SEM: Standard Error of Mean

**Table 6.** Percentage effect of mean difference of parameters of lipid profile for before and after atorvastatin and garlic extract administration.

| **Variables** | **Mean Difference** | **% Effect of before and after atorvastatin administration** | **Mean Difference** | **% Effect of before and after garlic administration** |
| --- | --- | --- | --- | --- |
| **TC (mmol/L)** | 1.28 | 41.6% | 1.30 | 43.2% |
| **HDL-C (mmol/L)** | 0.02 | 2.0% | 0.17 | 16.7% |
| **LDL-C (mmol/L)** | 0.72 | 47.4% | 0.68 | 44.7% |
| **TG (mmol/L)** | 0.15 | 28.3% | 0.13 | 24.4% |
| **CR** | 1.17 | 37.7% | 0.90 | 29.0% |

TC: Total cholesterol, TG: Triglyceride, LDL: Low-density lipoprotein, HDL: High-density lipoproteins, CR: Coronary Risk

**Table 7.**
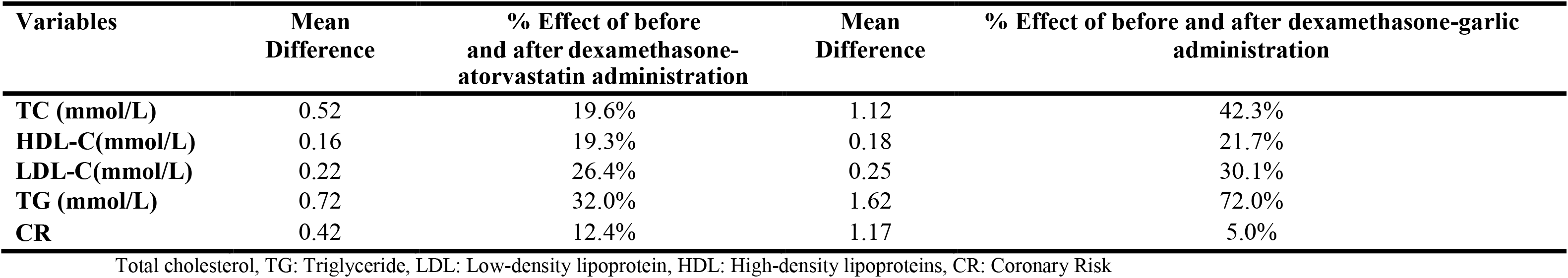
Percentage effect of mean difference of lipid profile for before and after dexamethasone-atorvastatin and dexamethasone-garlic extract.

| **Variables** | **Mean Difference** | **% Effect of before and after dexamethasone-atorvastatin administration** | **Mean Difference** | **% Effect of before and after dexamethasone-garlic administration** |
| --- | --- | --- | --- | --- |
| **TC (mmol/L)** | 0.52 | 19.6% | 1.12 | 42.3% |
| **HDL-C (mmol/L)** | 0.16 | 19.3% | 0.18 | 21.7% |
| **LDL-C (mmol/L)** | 0.22 | 26.4% | 0.25 | 30.1% |
| **TG (mmol/L)** | 0.72 | 32.0% | 1.62 | 72.0% |
| **CR** | 0.42 | 12.4% | 1.17 | 5.0% |

Total cholesterol, TG: Triglyceride, LDL: Low-density lipoprotein, HDL: High-density lipoproteins, CR: Coronary Risk

## DISCUSSION

### The effect of butter and dexamethasone on blood lipids

Hyperlipidaemia induction with butter and dexamethasone had an impact on the lipid profile parameters of the rats in the butter group as well as the dexamethasone group (Table 1). A similar effect was exerted in the lipid profiles of the rats as was reported by Adams and Talwalkar [27] who intimated from their study that giving rats 25% butter-enriched diet for 15 days will induce hyperlipidaemia. Also the results obtained from Table 1 agrees with the findings of Sowmya and Ananthi [28] who in their study; ‘Hypolipidaemic activity of *Mimosa pudica Linn* on Butter Induced Hyperlipidaemia in Rats’ reported that the administration of 400mg of butter/kg body weight in 10ml of buffered saline induced hyperlipidaemia in experimental rats.

Rats induced with hyperlipidaemia using dexamethasone (Dexamethasone Control Group) also showed a significant level of impact in their observed lipid profile parameters. When compared to the normal control group (NCG), the lipid parameters of the dexamethasone control group (DCG) being TC, LDL-C and TG showed a remarkable increase in their analysed values whilst HDL-C decreased. This result is in line with the outcome of the study done by Mahendran and Shyamala [29] and that of where the administration of 10mg/kg body weight of dexamethasone induced hyperlipidaemia in experimental rats.

Comparing coronary risk ratios of the butter control and the dexamethasone control groups to the normal control group showed an influence on the lipid profile parameters of the rats. The coronary risk ratio of the butter control group and the dexamethasone control group were higher compared to that of the normal group (Table 1). This study proved that, there was induction in both groups and this significantly influenced hyperlipidaemia development; a reason for the high coronary risk ratios.

### The effect of atorvastatin/garlic extract on hyperlipidaemia

Treatment of the hyperlipidaemic groups with Atorvastatin and aqueous garlic extract had an immense impact on the serum lipid and lipoprotein levels of both the butter-induced hyperlipidaemic rats and the dexamethasone-induced hyperlipidaemic rats. This study indicates that garlic reduces total cholesterol and LDL-cholesterol. Adler and Holub [30], reported similar observations in their study where there was, 11.5 % decrease of total cholesterol and 14.2 % decrease of LDL-cholesterol.

Treatment of the butter-induced hyperlipidaemia group with Atorvastatin (BAG) (Table 2) showed a moderately raised HDL-C level when compared to the HDL-C levels of the BCG and the NCG. It was also observed from the results of the Butter-Atorvastatin group (BAG) that the other serum lipoproteins; being TC and LDL-C reduced in their concentration levels in comparison to that of the BCG and NCG except for the TG levels that was slightly higher than that of the BCG and NCG. The result indicates that Atorvastatin is a standard positive control drug significantly influenced the blood lipid levels. This development is consistent with the study findings of Sheetal et al. [31]where Atorvastatin influenced the lipid and lipoprotein levels in a similar manner.

From the data obtained (Table 2), garlic extract had significant effects on theexperimental rats induced with hyperlipidaemia using butter enriched diet. There was an increase in the concentration of HDL-C and a corresponding reduction in the levels of TC and LDL-C except for TGs which was significantly raised when compared to that of the BCG and NCG. With the exception of the rise in TG level, this development is similar to the study findings of Ugwu and Omale [32].

### Effect of Atorvastatin on Dexamethasone-Induced Hyperlipidaemia

The co-administration of dexamethasone and atorvastatin showed a competitive effect on the lipid profile parameters of the rats due to their interaction. The HDL-C concentration upon the co-administration of atorvastatin and dexamethasone (as seen in the DAG group, Table 3) was observed to have decreased considering that of DCG and NCG. Also there was reduction in the concentration of TC in DAG group as against the concentration of TC in the DCG and NCG groups. These changes could be as result of the counter reaction of dexamethasone. Thus the drug was able to reduce the bloodstream levels of atorvastatin thereby making it less effective in treating the induced hypercholesterolaemia. This outcome was inconsistent with the findings of Chakraborty et al. [33] where Atorvastatin reduced the lipid levels of Triton X100 drug–induced hyperlipidaemia. It was also observed to be different from the outcome of another study by Pragda et al. [23] where the statin drug, Gemfibrozil was able to correct hyperlipidaemia induced by dexamethasone. The findings of this study therefore indicate that, the interaction of atorvastatin and dexamethasone had a competitive impact on the lipid levels of the experimental rats.

### Effect of Garlic Extract on Dexamethasone-Induced Hyperlipidaemia

The intra-gastric administration of 400mg/kg body weight of aqueous garlic extract on dexamethasone-induced hyperlipidaemic rats showed (as seen in Table 3) that garlic has bioactive lipid-lowering effects [30]. The LDL-C, TC and TG concentrations levels of DCG decreased significantly on treating the hperlipidaemic rats with aqueous garlic extracts. This implies that aqueous garlic extract had an effective anti-hyperlipidaemic impact on dexamethasone-induced hyperlipidaemia. This outcome confirms the study of [32] where aqueous garlic extract proved to be an effective lipid lowering agent. The outcome of this particular research also implicates aqueous garlic extract as anti-hyperlipidaemic agent thereby confirming the findings of the researchers; [34-36] who credited garlic extract as an effective hypolipidaemic agent.

### The Impact of Garlic Extract on Coronary Risk

The administration of aqueous garlic extract on butter-induced hyperlipidaemic rats showed significant decline in the coronary risk ration compared to that of BCG and NCG. Thus, treatment of hyperlipidaemic rats with aqueous garlic extract decreased the coronary risk ratio. This implies that rats treated with aqueous garlic extract had lower risk of getting coronary heart or cardiovascular related diseases. Thus the garlic was effective in conferring cardio protection on the rats thereby making them less prone to coronary heart diseases [30]. The garlic extract exerts similar bioactive effects like the atorvastatin; thereby reducing the risk rate at which the experimental rats are exposed to coronary heart diseases with respect to their lipid levels. Similar observations were reported by [30]. After an intra-gastric administration of the hyperlipidaemic rats with aqueous garlic extract, it was observed that the mean body weight increased from week 1 through week 3, 4 but no statistical significance was observed, however, a significant increase from week 4 to week 5 (p=0.048). The value suggests that garlic extract had no reduction influence on the body weight of the experimental rats.

The study concludes that butter and dexamethasone were effective in inducing hyperlipidaemia in the experimental rats. Administering dexamethasone through subcutaneous injection increased the lipid profile parameters of the rats. Thus, dexamethasone successfully induced hyperlipidaemia in the experimental rats. Also administering aqueous garlic extract on the butter-induced hyperlipidaemic rat affected the lipid parameters. The aqueous garlic extract increased the concentration of HDL-C and simultaneously decreased the concentration levels of LDL-C, TC and TG in the butter-induced hyperlipidaemic rats. The aqueous garlic extract had a lipid-lowering effect on dexamethasone-induced hyperlipidaemia and it exhibited a similar effect as the standard drug, Atorvastatin. The extract was effective in reducing the coronary risk ratio of the hyperlipidaemic rats to near normal; an indication of the extract having cardio-protective effects. Comparatively, the aqueous garlic extract had less effect on the body weight of butter-induced hyperlipidaemic rats to that of the Atorvastatin. Based on the outcome of this research study, a further study into the subject matter with an inclusion of the following suggested ideas will contribute to biomedical research knowledge as well as help inform choices when it comes to choosing garlic as an alternative treatment for hyperlipidaemic conditions.

Limitations of study include study was subject to some degree of sampling or selection bias. In addition, information from experimental rats’ exposure is subject to observational bias and inability to determine the incidence or prevalence of a condition because experimental rats are collected by purposive sampling.

## Acknowledgements

The authors appreciate the support from Prof. F.A. Yeboah, Dr. Christian Obirikorang, Prof. Robert Ngala as well as Mr. Thomas Ansah for their academic support and proof reading the manuscript.

## Availability of data materials

The datasets generated and/or analyzed during the current study are not publicly available but are available from the corresponding author on reasonable request.

## Authorship Contribution

GS was responsible for the conception, design, participants’ recruitment and collection of present study. GS, EA, MS were involved in the data analysis. EA, GS and RPA drafted the article. All the authors proofread and approved the final version for publication.

